# Systematic delineation of signaling and epigenomic mechanisms underlying microglia inflammatory activity in acute and chronic brain pathologies

**DOI:** 10.1101/2022.08.04.502805

**Authors:** Andre Machado Xavier, Félix Distéfano-Gagné, Nesrine Belhamiti, Sarah Belhocine, Sara Bitarafan, Alexia Falle, S. Fiola, Serge Rivest, David Gosselin

## Abstract

Microglia promptly mount an inflammatory response following detection of infectious agents or injuries in the central nervous system. Such function fundamentally depends upon dynamic modulation of gene expression. However, the signaling and epigenomic mechanisms that regulate the transcriptional process underlying microglial inflammatory activity are not well understood. To address this, we used RNA-seq, ChIP-seq and ATAC-seq to delineate gene signatures and activity across the repertoire of genomic regulatory elements of microglia engaged in acute and chronic neuroinflammatory activity. Systematic interrogations of the microglial population over time during a systemic inflammatory response revealed a coordinated, sequential activation of multiple gene programs associated with defense response, translation and cell cycling. Activation of these programs occurred in parallel with gain and loss of activity at 4,080 and 3,119 genomic cis-regulatory elements, respectively. Furthermore, computational analyses identified key transcriptional regulators, including Ets, AP-1, C/epb, Nf-κB, Irf, Runx, c-Myc and E2f family members, that display differential propensity for activity at gene promoters and promoter-distal cis-regulatory elements. Gene expression analyses also suggested that the transcriptional process likely contribute to the effective activity of numerous transcriptional regulators through the modulation of their mRNA levels. Finally, characterization of CD11c-positive microglia that emerge with chronic demyelinating brain lesions suggested that Egr2, Mef2 members and E-box-binding factors such as Tfeb and Mitf contribute to the enhanced phagosomal activity of this inflammatory subset. Loss-of-function experiments validated that Mef2a in microglia is necessary for the acquisition of the CD11c-positive phenotype. Collectively, these results demonstrate that the inflammatory activity of microglia arises through an intricate, ultimately context-dependent, interplay between signaling pathways, genomic regulatory elements and the transcriptional machinery.

## Introduction

As tissue-resident macrophages, microglia are essential effectors of inflammatory responses in the CNS, or neuroinflammation(1). These responses occur upon detection of infections and injuries, but also in chronic neurodegenerative disorders such as Alzheimer’s disease (AD) and multiple sclerosis (MS)(2, 3). When properly executed, inflammatory responses efficiently eliminate pathogens, limit damage following injuries, promote repair and resolve themselves in an orderly manner(4). In chronic neurodegenerative disorders however, and for reasons that are not well understood, microglia engage in sustained inflammatory activity that can exacerbate damage to the neural tissue(2, 5). These characteristics highlight the complex and varied nature of neuroinflammatory responses, while also suggesting intricate, context-dependent regulatory mechanisms.

The inflammatory cell functions of microglia arise through transcriptional induction of gene coding for various molecular mediators of inflammation, including cytokines, chemokines and proteases(6). Moreover, while inflammatory microglia associated with different lesions express many genes in common, the overall signature is context-specific(7). Indeed, systematic comparisons of microglia isolated from mouse models of AD, MS and amyotrophic lateral sclerosis (ALS) revealed common upregulation of multiple inflammatory/immune genes, including *Apoe, Clec7a, Cst7*, albeit to varying levels across pathologies(8, 9). Alternatively, acute activation of mouse microglia triggered during a systemic inflammatory response led to specific induction of *Map3k8* and *Socs3* as compared to inflammatory microglia associated with aging, AD, or ALS(10). Such context-dependent responses presumably ensure that microglial inflammatory activity operates in accordance with the physiological demands of the surrounding, perturbed, brain microenvironment.

Induction of inflammatory gene programs in macrophages depends upon coordinated stimulation of multiple signaling pathways that ultimately trigger activation of potent pro-inflammatory transcription factors (TFs), including Nf-κB, AP-1, Irf and Stat family members(11). Efficient delivery of regulatory input by these factors to the transcriptional machinery requires calibrated interactions between TFs and chromatin(12). These interactions take place at genomic cis-regulatory elements, including gene promoters and promoter-distal elements. Promoter regions support the full assembly of the RNA polymerase II complex required for effective gene transcription. In contrast, promoter-distal elements provide the majority of binding sites for transcription factors active in a cell; their number indeed supersedes that of poised or active promoters(13). As such, promoter-distal elements help calibrate context and burst frequencies of transcription of their target genes(14, 15). Coherent with this, early work in primary macrophages revealed that 90% of chromatin binding events of the p65 Nf-κB subunit following Tlr4 stimulation occurs at promoter-distal elements(16). More recently, a study reported gains in nucleosome histone acetylation, an epigenetic marker for elevated local transcription factor binding activity(17, 18), at 4,201 promoter-distal elements in Kupffer cells in the lesioned liver compared to healthy tissue(19). Notably, these elements displayed enrichment for DNA motifs recognized by Runx, Egr2, and bZip factors AP-1 and Atf family members, which suggests that these transcription factors are highly active in these macrophages in lesion circumstances.

To date, multiple signaling and transcriptional regulators have been linked to regulation of inflammatory activity in the brain. For example, IκB kinase (IKK) drives Nf-κB activity in microglia in mouse models of AD and Irf5 regulates microglial activity following strokes(20, 21). However, myriads of pro-inflammatory signaling pathways are co-activated in conditions of brain lesions, but how these signals integrate with one another and how they interact with regulatory elements in microglia is not well understood. To this end, there is evidence suggesting that the microglial repertoire of regulatory elements is highly responsive to disturbances in signaling input received by microglia.

For example, removing microglia from the brain and transferring them to tissue culture conditions results in the loss of activity at 6,382 promoter-distal regulatory elements(22). Notably, these regions exhibited enrichment for DNA motifs linked to Smad and Mef2 transcription factors, among others, which is coherent with the role of these factors in specifying microglial cell functions in the healthy brain.

Here, we sought to understand how inflammatory pathways and transcription factors coordinate transcription on genome-wide level in microglia during brain perturbations. For this, we characterized the epigenomic and transcriptional response of microglia engaged in heighten immune activities associated with different neuroinflammatory conditions. Globally, our results suggest that microglial inflammatory activity arises from activation and suppression of combinations of common and context-specific genomic regulatory elements.

### Systemic injection of lipopolysaccharide elicits dynamic, time-dependent transcriptional responses in microglia

We first sought to understand how microglia transition from a state of homeostatic activity to an inflammatory state at the transcriptional level. For this, we profiled gene expression over time in microglia isolated from mice undergoing a systemic inflammatory response triggered by lipopolysaccharide (LPS). Injection of LPS (1 mg/kg) in the peritoneal cavity of adult mice elicits a highly reproducible, multi-phase inflammatory response throughout the body, including the brain(23). Hence this model allows for the precise characterization of changes in microglial gene expression across time. We sacrificed C57BL/6 mice 3, 6, 24 hours (h) and 7 days following LPS injection and isolated microglia using fluorescence-activated cell sorting (FACS). We defined microglia as live, CD45^Low^CD44^Low^CD11b^+^ single cells (Fig.S1A). We also assessed CD11c expression to monitor for possible emergence of distinct inflammatory microglial subsets(24). Overall, the microglial population as identified by expression of the cell surface proteins remained homogenous and similar over the time points interrogated (Fig.S1B, C). Yet, principal component (PC) analysis of the gene profiles of these microglia, as assessed by RNA-seq, revealed that marked alterations at the transcriptional level occurred in a time-dependent manner, and particularly over the first 24 hours (Fig.1a; Table S1). Indeed, compared to non-treated mice, microglia at the 3, 6 and 24h time points upregulated significantly 822, 734, 267 genes, respectively (Fig. 1B; log2 fold change (lfc) ≥ 0.58, false discovery rate (FDR)-adjusted p-value ≤ 0.01). Concomitantly, 350, 667, and 157 genes were downregulated, again respectively. This substantial overhaul in the transcriptional output of microglia however was largely restricted to the early time points; indeed, by day 7, microglia had largely returned to a transcription ground state.

**Figure 1:**
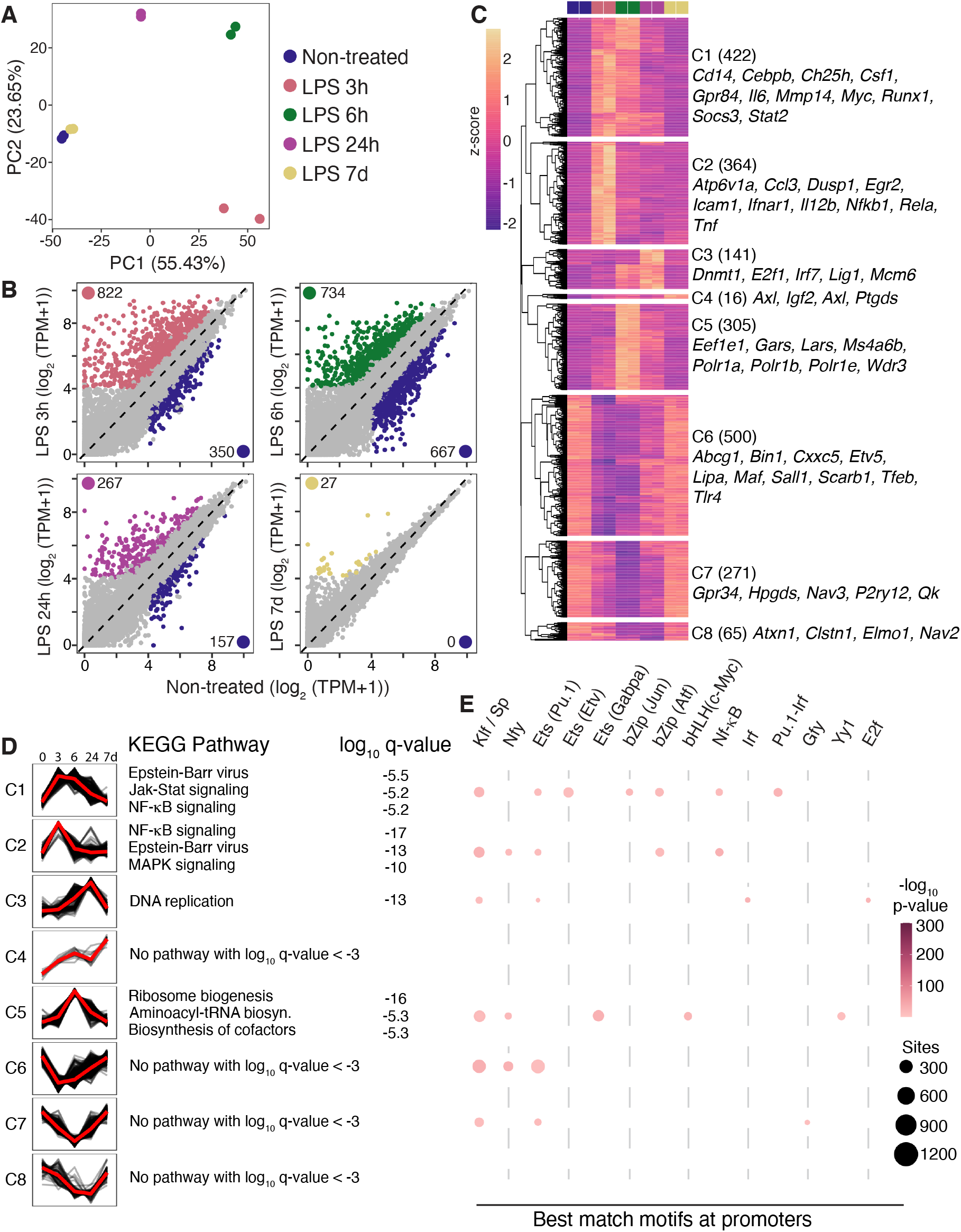
Temporal modulation of gene expression in microglia triggered by systemic injection of LPS. **(A)** Principal component analysis of RNA-seq data from microglia isolated from the brain of mice at different time points after LPS injection. **(B)** Scatterplots of RNA-seq data of microglia isolated at various time points after LPS injection, compared to non-treated mice. Differentially expressed genes identified by DESeq2 (log2 fold-change (FC) ≥ 0.58, FDR-adjusted p-value ≤ 0.01) are colored-coded. **(C)** Hierarchal clustering of significantly modulated genes identified in B. **(D)** Summary profiles and KEGG pathways enrichment analysis of gene clusters identified. **(E)** Table summarizing results from de novo DNA motifs analyses performed on promoters of gene clusters identified.

To achieve a more precise understanding of the different gene programs modulated in microglia and their temporal dynamics of expression, we performed hierarchal gene clustering based on all the differentially expressed genes identified above. This revealed 8 main temporal profiles of expression, with 5 that exhibited distinct patterns of induction and 3 that comprised downregulated genes (Fig. 1C, D). Of note, dominant profiles C1 and C2 included genes markedly induced as early as 3h. Expression plateau dynamics discriminated these clusters. Indeed, while induction of C1 genes remained elevated until at least 6h, that of C2 genes was very transient, peaking at 3h. Globally, both clusters encoded genes involved in antimicrobial activity (e.g., *Cd14, Ch25h, Il6, Tnf*), and regulation of Nf-κB, Jak-Stat, and MAP kinase pathways (e.g., *Socs3, Stat2, Rela, Dusp1*). A burst of transcriptional induction then followed at 6h (i.e., C5 genes) and these comprised molecular effectors relevant to protein synthesis such as *Eef1e1, Gars, Polr1a, Polr1b*. Finally, C3 genes peaked in expression at the 24h mark and these were collectively strongly associated with DNA replication. While the induction of genes was often defined by strong association with defined cellular processes, such associations were largely absent for downregulated genes. Nevertheless, we noted decreased expression of numerous genes involved in the regulation of microglial activity in the healthy brain, including *Bin1, Maf, P2ry12, Sall1* and *Tfeb*. Together, these data indicate that multiple waves of functionally distinct transcriptional programs are sequentially triggered in microglia during the first 24 hours of an LPS-induced systemic inflammatory response. We note that similar observations were previous made, albeit in a more severe model that used an LPS dose of 2.5 mg/kg(25).

We next investigated whether the identified profiles may be regulated by distinct combinations of TFs acting at gene promoters. To achieve this, we performed, for each of the profiles, DNA motif analyses centered on areas of accessible chromatin, defined by ATAC-seq, located within the promoter regions of their respective genes. Overall, all clusters except minor clusters C4 and C8 displayed combinations of enrichment for general promoter motifs linked to Klf/Sp, Nfy, and Ets TFs (Fig. 1E). Although less prominent, we also observed evidence for more cluster-biased enrichment. For example, some C1 and C2 gene promoters both contained potential binding sites for bZip and Nf-κB factors. There was also minor enrichment for the Myc motif at C5 genes promoters and for the E2f motif at those of genes that peaked at 24h (i.e., C3). Alternatively, promoters of downregulated clusters displayed no major enrichment for motifs beyond those recognized by Klf/Sp, Nfy, and Ets family members. Therefore, a combination of input, albeit relatively weak, provided by broadly active general TFs and more context-dependent pathways may regulate activity at promoter elements in microglia during a systemic inflammatory episode.

### Transient alterations in histone acetylation in inflammatory microglia activated during a systemic inflammatory response

Along with gene promoter regions, promoter-distal genomic cis-regulatory elements (CREs) play a key role in coordinating transcription; TF activity at these discrete regions contributes specification of timing and spatial parameters of gene transcription(11, 15). To gain insights into how these elements may help coordinate transcription in microglia as they mount an inflammatory response, we assessed their activity levels following systemic injection of LPS. For this, we profiled with ChIP-seq abundance of the H3K27ac epigenetic mark, a marker of transcriptional activity at genomic regulatory elements(17, 18), on a genome-wide level in microglia isolated at 3, 6, 24 h and 7 days after injection of LPS. Principal component (PC) analysis revealed a strong partitioning on PC1, which accounted 47.7% of data variance (Fig. 2A). Mechanisms linked to inflammatory activation likely drove PC1 as the early LPS time points clustered together on one extremity, and the non-treated and LPS 7d microglia together on the other. Pairwise comparisons with non-treated microglia corroborated the substantial alterations in promoter-distal H3K27ac over the first 24 hour of the LPS-associated response (Fig. 2B, C). Unlike the mRNA transcriptional response however, for which substantial changes occurred as early as 3h, alterations in H3K27ac were relatively delayed. Indeed, while only 1,068 sites displayed significant changes in H3K27ac at 3h, important modulations occurred by 6 and 24h, with 3,167 and 3,983 promoter-distal sites displaying differential H3K27ac levels, respectively (FDR-adjusted p-value ≤ 0.05, two-fold or greater tag difference; Fig. 2B). Hierarchal clustering based on regions that displayed differential H3K27ac in LPS-associated microglia revealed two dominant profiles of gains of H3K27ac in microglia in the LPS model; C1 included 1,536 regions that peaked in gains at the 6h timepoint and C3 comprised 1,069 that peaked at 24h (Fig. 2C, D). Regions that lost H3K27ac, i.e., C4, C5 and C6, also did so according to different dynamics.

**Figure 2:**
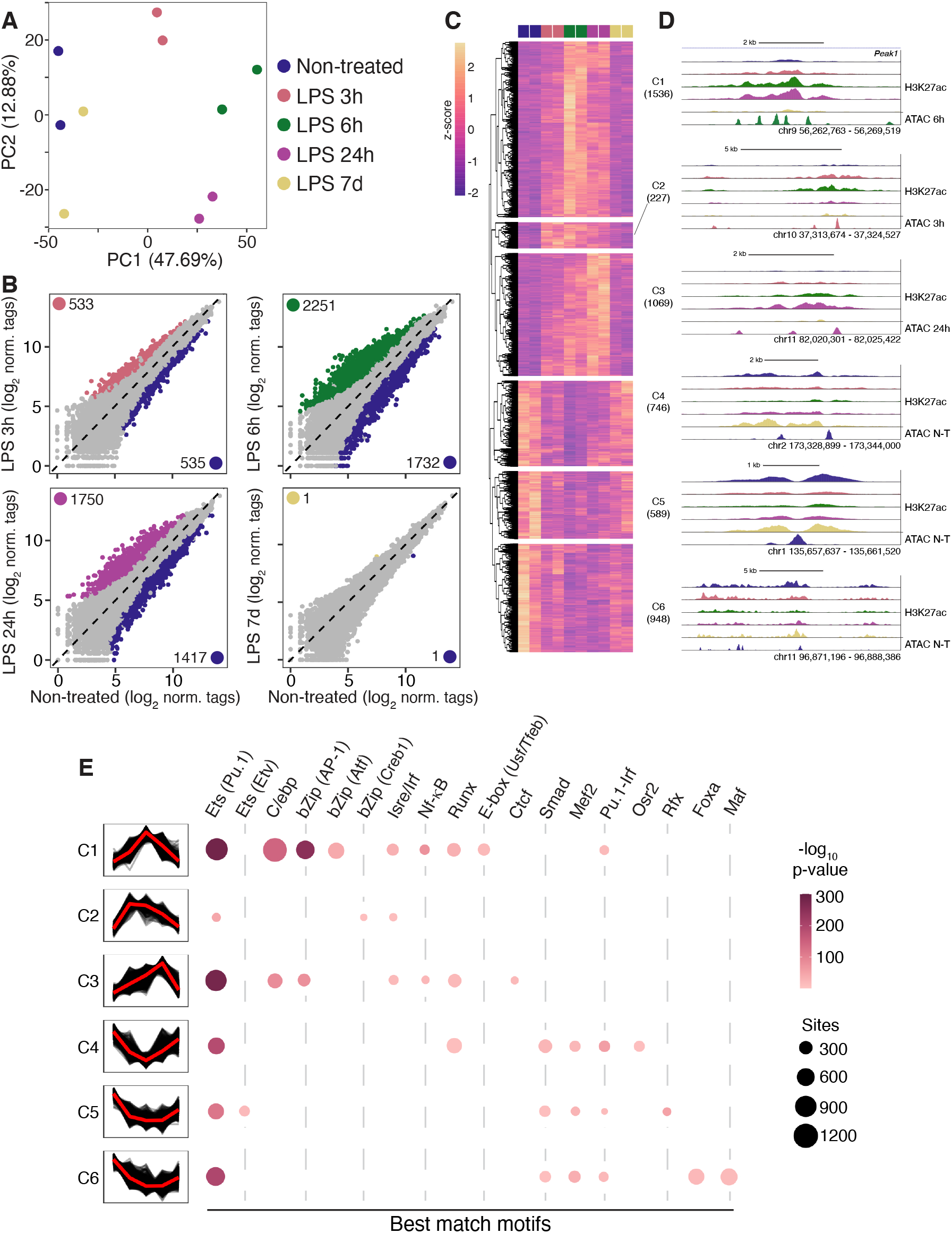
Characterization of H3K27ac deposition at promoter-distal cis-regulatory elements in microglia following systemic injection of LPS. **(A)**. Principal component analysis of H3K27ac ChIP-seq data at promoter-distal CREs of microglia isolated from the brain of mice at different time points after LPS injection. **(B)** Scatterplots of H3K27ac abundance at distal CREs in microglia isolated at various time points after LPS injection, compared to non-treated mice, as assessed by ChIP-seq. Differentially enriched CREs identified by DESeq2 (FDR-adjusted p-value ≤ 0.05, two-fold or greater tag difference) are colored coded. **(C)** Hierarchal clustering of significantly modulated CREs identified in B. **(D)** UCSC browser displays of H3K27ac data at loci of genomic elements representative of the clusters identified in C. ATAC-seq data associated with each cluster peak time are also appended. **(E)** Table summarizing results from de novo DNA motifs analyses performed on the clusters of distal CREs identified.

We next performed DNA motif enrichment analyses to identify potential TFs that may control activity at the different cluster of modulated distal CREs. Again, these were centered on ATAC-seq-defined areas accessible chromatin encompassed with the H3K27ac regions of interest. Results revealed a strong enrichment for Ets factor Pu.1 across all clusters, which is consistent with Pu.1 acting as a lineage-determining TF that sets up promoter-distal CREs in macrophages (Fig. 2E)(26). With respect to elements that specifically gain H3K27ac, and major clusters C1 and C3 in particular, bZip factors AP-1 and C/ebp, and to lesser extent by Isre/Irf, Nf-κB, Runx, and E-box motifs, dominated the motifs landscape. Analyses of regions that lost H3K27ac as part of the LPS-associated response revealed that these may be regulated by a large array of TFs typically associated with homeostatic activity of microglia. These included Smad, Mef2, Foxa and Maf. Finally, Pu.1-Irf composite sequence and Runx motifs were enriched at both upregulated and downregulated distal CREs, which may suggest context- and site-dependent activity for these factors.

Collectively, these results suggest that a complex interplay between multiple signaling pathways, TFs and promoter-distal CREs underlies the acquisition of heightened immune defense capabilities by microglia. These results are also coherent with a hierarchal model of macrophage activation, with AP-1 and C/epb providing dominant input, and Nf-κB and Irf, while key inflammatory factors, act in more restricted capacity(13). Lastly, these results also underlie that a general suppression of homeostatic input accompanies microglial inflammatory activation.

### Systemic inflammation induces a delayed burst of microglial proliferation

Previous studies reported that microglia proliferate as part of an LPS-induced systemic inflammatory response(27-29). The transcriptional induction of genes involved in DNA replication at the 24h time point also suggested that a cell cycle program is triggered in this model. Flow cytometry analyses of Ki67 expression, a marker of cell proliferation, confirmed that a large proportion of microglia proliferate over the first 72 hours after LPS injection (Fig. 3A, B).

**Figure 3.**
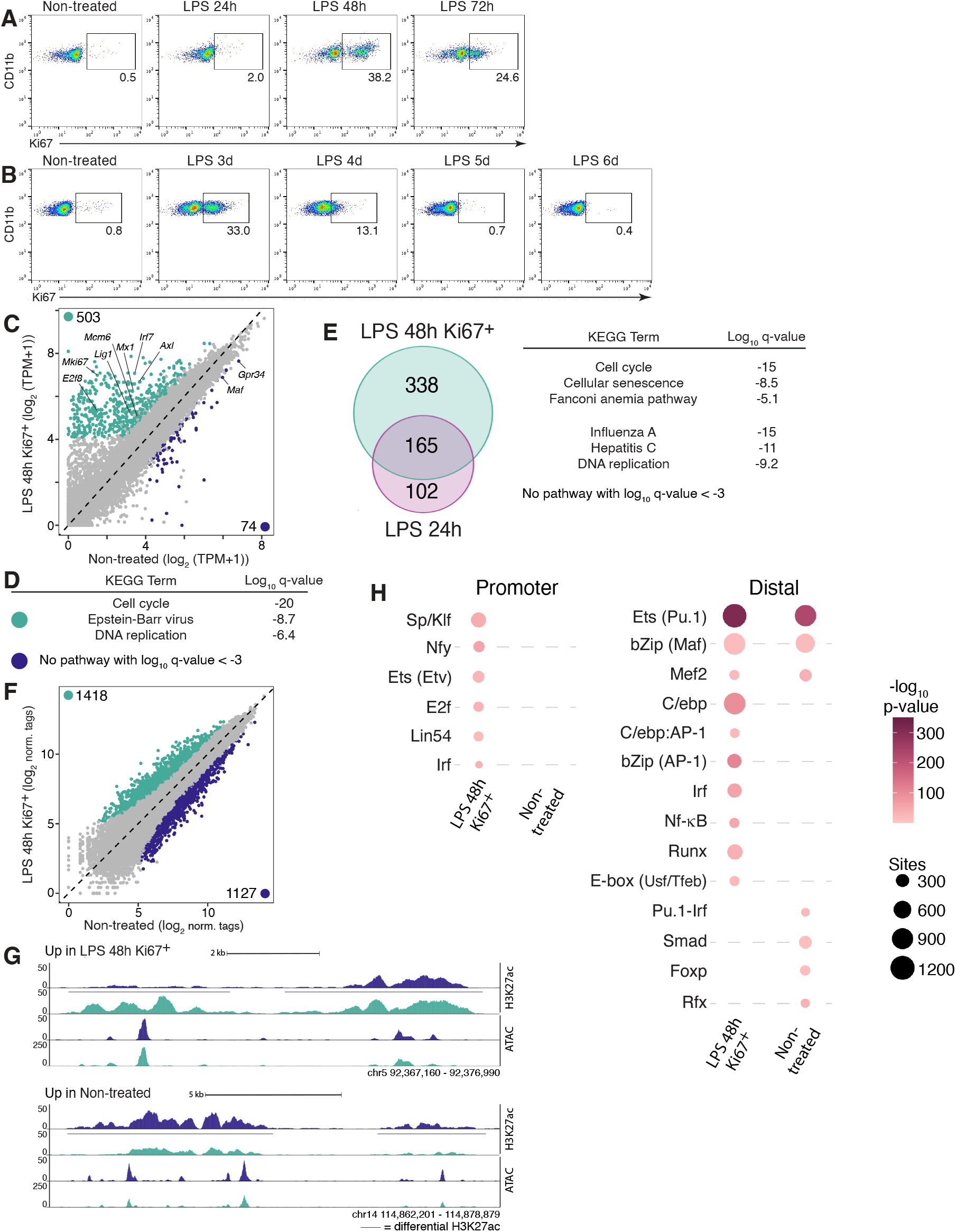
Transcriptional and epigenomic features of inflammatory, proliferative microglia induced following systemic injection of LPS. **(A)** Flow cytometry data depicting detection of Ki67^**+**^ microglia in the brain of mice between 24 and 72 h after LPS injection. **(B)** Flow cytometry data depicting detection of Ki67^**+**^ microglia in the brain of mice between 3 and 6 days after LPS injection. **(C)** Scatterplot of RNA-seq data of RFP^High^/Ki67^**+**^ microglia from LPS 48 h mice and quiescent RFP^Low^/Ki67^**−**^ microglia isolated from the brain non-treated mice. Differentially expressed genes identified by DESeq2 (log2 FC ≥ 0.58, FDR-adjusted p-value ≤ 0.01) are colored coded. **(D)** KEGG pathways analysis of genes significantly more highly expressed in LPS 48h Ki67^**+**^ vs quiescent Ki67^**−**^ microglia from non-treated mice, and vice versa. **(E)** Venn diagram depicting genes that are upregulated in both LPS 48h Ki67^**+**^ and LPS 24h microglia compared to non-treated, quiescent Ki67^**−**^ microglia, and associated KEGG pathway analysis. **(F)** Scatterplot of H3K27ac abundance at distal CREs, comparing LPS 48 h Ki67^**+**^ vs non-treated/quiescent Ki67^**−**^, as assessed by ChIP-seq. Differentially enriched CREs identified by DESeq2 (FDR-adjusted p-value ≤ 0.05, two-fold or greater tag difference) are colored coded. **(G)** UCSC browser displays of H3K27ac and ATAC-seq data at representative loci of genomic elements that exhibit significantly higher (top) or lower (bottom) levels of H3K27ac in LPS 48 h Ki67^**+**^ vs non-treated/quiescent Ki67^**−**^ microglia. **(H)** Table summarizing results from de novo DNA motifs analyses performed on differentially active promoters and distal CREs identified in C and F, respectively.

We next sought to better understand the transcriptional mechanism underlying microglial proliferation in this LPS model. For this, we first assessed the gene signature of Ki67^**+**^ proliferative microglia at 48h post-LPS injection using Ki67^RFP^ mouse line, in which the RFP signal correlates with abundance of the Ki67 protein(30); we selected the 48h time point because the proportion of Ki67^**+**^ microglia was maximal then (Fig. 3A). Compared to their RFP^Low^/Ki67^**−**^ counterparts, RFP^High^/Ki67^**+**^ microglia expressed at significant higher levels 101 genes, including *Ccnb2, Mki67* and *Plk1* (Fig. S2A; Table S2). KEGG pathway and gene ontology (GO) analyses confirmed that LPS 48h Ki67^**+**^ microglia display a prominent cell cycle gene signature, and that they are likely at the G2/M stage of the process, in agreement with a previous study (Fig.S2B)(31). Also coherent with this previous work, profiles H3K27ac across the repertoire of promoter-distal CREs were highly similar between the two subsets (Fig. S2C).

The Ki67^**+**^ vs Ki67^**−**^ RNA-seq analysis above did not reveal expression differences for gene linked to immune defense response. This may either reflect a complete return of expression of these immune genes to a basal state by 48h or instead a specific gain of transcription limited to cell cycle gene in Ki67^**+**^ microglia. Thus, to further examine the immune profile of LPS-associated proliferative microglia, we compared LPS-Ki67^**+**^ subset to non-treated, Ki67^**−**^ microglia from the heathy adult brain. Overall, a total of 577 genes were differentially expressed between these two subsets (Fig. 3C; Table S2). Notably, among the 503 genes upregulated in LPS 48h Ki67^**+**^ microglia, there was a strong enrichment for genes linked to antiviral immunity, including *Axl, Irf7*, and *Mx1*, and cell cycling, including *Lig1, Mcm6* and *Mki67* (Fig. 3C). Furthermore, 62% (165/267) of the genes upregulated in microglia at 24 h vs non-treated condition retained elevated expression in LPS 48h Ki67^**+**^ microglia and these were also strongly associated with antiviral gene programs (Fig. 3E). Thus, proliferating microglia in the LPS model conserve a robust inflammatory gene signature. In contrast, 74 genes were downregulated in LPS 48h Ki67^**+**^ microglia compared to the non-treated condition (Fig. 3C). However, KEGG pathway analysis did not reveal strong association for these with a catalogued biological process (Fig. 3D). Finally, the epigenome of LPS 48h Ki67^**+**^ microglia also displayed a pro-inflammatory signaling signature (Fig. F to H). Indeed, CREs in this subset that had higher levels of H3K27ac compared to non-treated microglia were also enriched for AP-1, C/ebp, Irf, Nf-κB and Runx motifs. Of interest, these were not enriched at promoters of upregulated genes in LPS 48h Ki67^**+**^ microglia. Instead, these displayed enrichment for E2f and Lin54, consistent with previous work on proliferative microglia(31). Together, these results support distinct mechanisms of gene regulation necessary for microglial cell proliferation and inflammatory activity. While the former may primarily rely upon sets of TFs acting preferentially at promoters, the latter appear to be proportionally more dependent upon different sets of TFs acting at CREs.

We and others previously showed that microglia that rapidly proliferate on a large scale to repopulate the brain following depletion by treatment with Csf1r inhibitor PLX5622 also transcribe inflammatory genes at high levels(31, 32). Therefore, we next sought to determine whether there is a convergence on an overall gene signature for microglia that proliferate massively and rapidly, albeit in different contexts. For this, we first compared with principal component analysis the transcriptomes of microglia that proliferate in different settings, including during brain development on post-natal day 8 (PND8), 3 days after termination of PLX5622 treatment, and the LPS 48h time point. Principal component analysis confirmed high similarities between LPS 48h Ki67^**+**^ and PLX-Ki67^**+**^ microglia, which in turn suggests a pro-inflammatory profiles to these subsets (Fig. 4A). Differential gene expression analyses corroborated this; LPS 48h Ki67^**+**^ expressed at significant higher and lower levels only 74 and 22 genes, respectively (Fig. 4B; Table S2). Of interest, while many upregulated genes in the LPS model intersected with regulation of TNF signaling, those more highly expressed in the PLX model, such as *Dhcr24* and *Sqle*, were linked to sterol biosynthesis (Fig. 4C). This may indicate that the availability and source of cholesterol required for microglial proliferation is highly subject to context-associated parameters.

**Figure 4.**
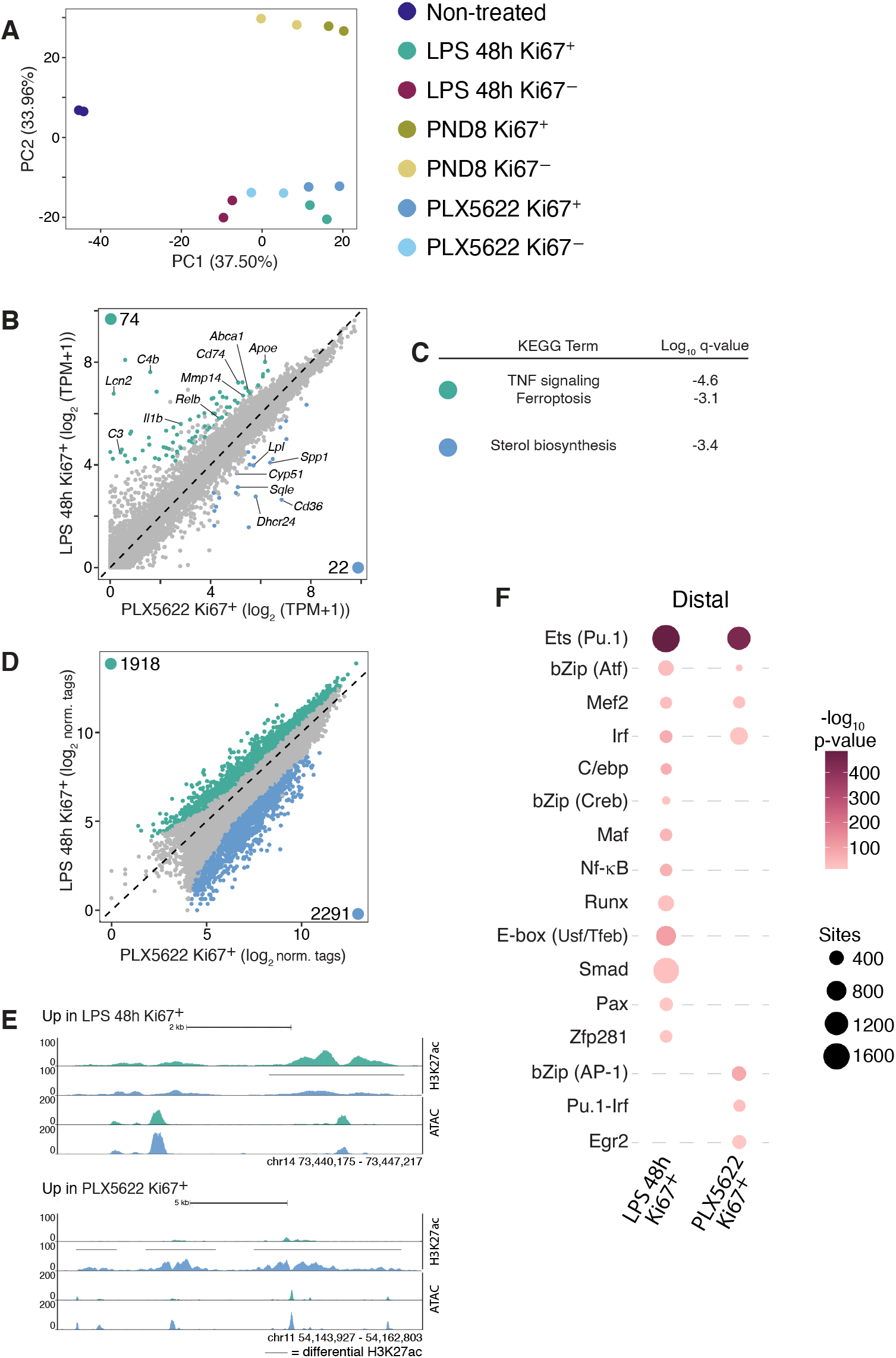
Microglia that abruptly proliferate on a large-scale display similar gene signature independent of physiological context. **(A)** Principal component analysis of RNA-seq data from RFP^Low^/Ki67^**−**^ and RFP^High^/Ki67^**+**^ microglia isolated from different physiological circumstances. **(B)** Scatterplot of RNA-seq data of RFP^High^/Ki67^**+**^ microglia from LPS 48 h mice and RFP^High^/Ki67^**+**^ microglia isolated from mice 3 days after termination of 17 days of PLX5622 diet. Differentially expressed genes identified by DESeq2 (log2 FC ≥ 0.58, FDR-adjusted p-value ≤ 0.01) are colored coded. **(C)** KEGG pathways analysis of significantly differently expressed genes identified in B. **(D)** Scatterplot of H3K27ac abundance at distal CREs, comparing LPS 48 h Ki67^**+**^ vs PLX5622 Ki67^**+**^, as assessed by ChIP-seq. Differentially enriched CREs identified by DESeq2 (FDR-adjusted p-value ≤ 0.05, two-fold or greater tag difference) are colored coded. **(E)** UCSC browser displays of H3K27ac and ATAC-seq data at representative loci of genomic elements that exhibit significantly higher (top) or lower (bottom) levels of H3K27ac in LPS 48 h Ki67^**+**^ vs PLX5622 Ki67^**+**^ microglia. **(F)** Table summarizing results from de novo DNA motifs analyses performed on differentially active promoters and CREs identified in B and D, respectively.

The high similarities between the LPS 48h Ki67^**+**^ and PLX Ki67^**+**^ gene signatures could also suggest concordance for their associated profile of signaling activity at promoter-distal CREs. Contrary to this prediction, direct comparisons of H3K27ac ChIP-seq data revealed highly divergent profiles, with 1918 and 2291 elements being respectively more and less acetylated in LPS 48h Ki67^**+**^ compared to PLX Ki67^**+**^ (Fig. 4D, E). Of interest, motif analysis also suggested that different configurations of signaling pathways operate in these different subsets (Fig. 4F). For example, while motifs linked to C/ebp, E-box, Nf-κB, and Atf were more prominent in LPS 48h Ki67^**+**^ microglia, the PLX subset was characterized by relatively elevated AP-1, Pu.1-Irf8 and Egr2 signatures.

Together, these data provide further evidence for a robust coupling between inflammatory gene programs and the ability of microglia to proliferate rapidly on a large-scale. However, they also indicate that while inflammatory microglia that proliferate in different physiological context may converge on a similar gene signature, the underlying regulatory landscape may differ substantially.

### Transcriptional mechanisms underlying heightened phagocytic activity of microglia in chronic demyelinating lesions

The inflammatory transcriptional activity microglia triggered in the systemic LPS model is very dynamic and transient, with microglia largely returning to a basal state by the seventh day after injection. Thus, the extent to which underlying regulatory mechanisms of transcription are relevant to inflammatory microglia associated with more chronic pathologies is not evident. To address this, we investigated gene regulation in CD11c^**+**^ microglia that accumulate with chronic demyelinating lesions triggered in mice by cuprizone (CPZ) neurotoxin. In this model, ingestion of CPZ induces substantial damage to oligodendrocytes throughout the brain that is accompanied by a strong inflammatory response(33). We isolated CD11c^**+**^ and CD11c^**−**^ microglia subsets after 4 weeks of CPZ and first performed RNA-seq (Fig. 5A). Overall, these subsets displayed relatively similar gene signature, with only 177 genes being differentially expressed between the two subsets (Fig. 5B; Table S3). Among others, the CD11c^**+**^ subset transcribed *Igf1* mRNA at higher levels, which is coherent with published literature on the CD11c^**+**^ microglial phenotype(34, 35). Comparison of the CD11c^**+**^ subset to CD11c^**−**^ microglia isolated from the healthy adult brain predictably yielded more pronounced differences (Fig. 5C; Table S3). Indeed, 479 and 197 genes were significantly upregulated and downregulated, respectively, in CD11c^**+**^ microglia. Notably, gene programs linked to phagosomal and antiviral immune activity, and cholesterol metabolism characterized the upregulated transcriptional signature of the CD11c^**+**^ microglia subset. Genes encoding effector of these functions included *Apoe, Abcg1, Ch25h*, and *Lyz*. Lastly, substantial alterations in activity levels of promoter-distal CREs also occurred in CD11c^**+**^ microglia in the CPZ model (Fig. 5D, E; Fig.S3A, B). For example, 4501 elements gained H3K27ac in CD11c^**+**^ microglia compared to basal microglia, and these displayed strong enrichment for Pu.1-Irf8, Egr2, E-box (e.g., Tfeb/Usf) in addition to robust Pu.1, AP-1 and C/epb signatures (Fig. 5D to F). Overall, these results depict the CD11c^**+**^ subset as capable of potent immune activity combined with high phagocytic, cargo elimination potential.

**Figure 5.**
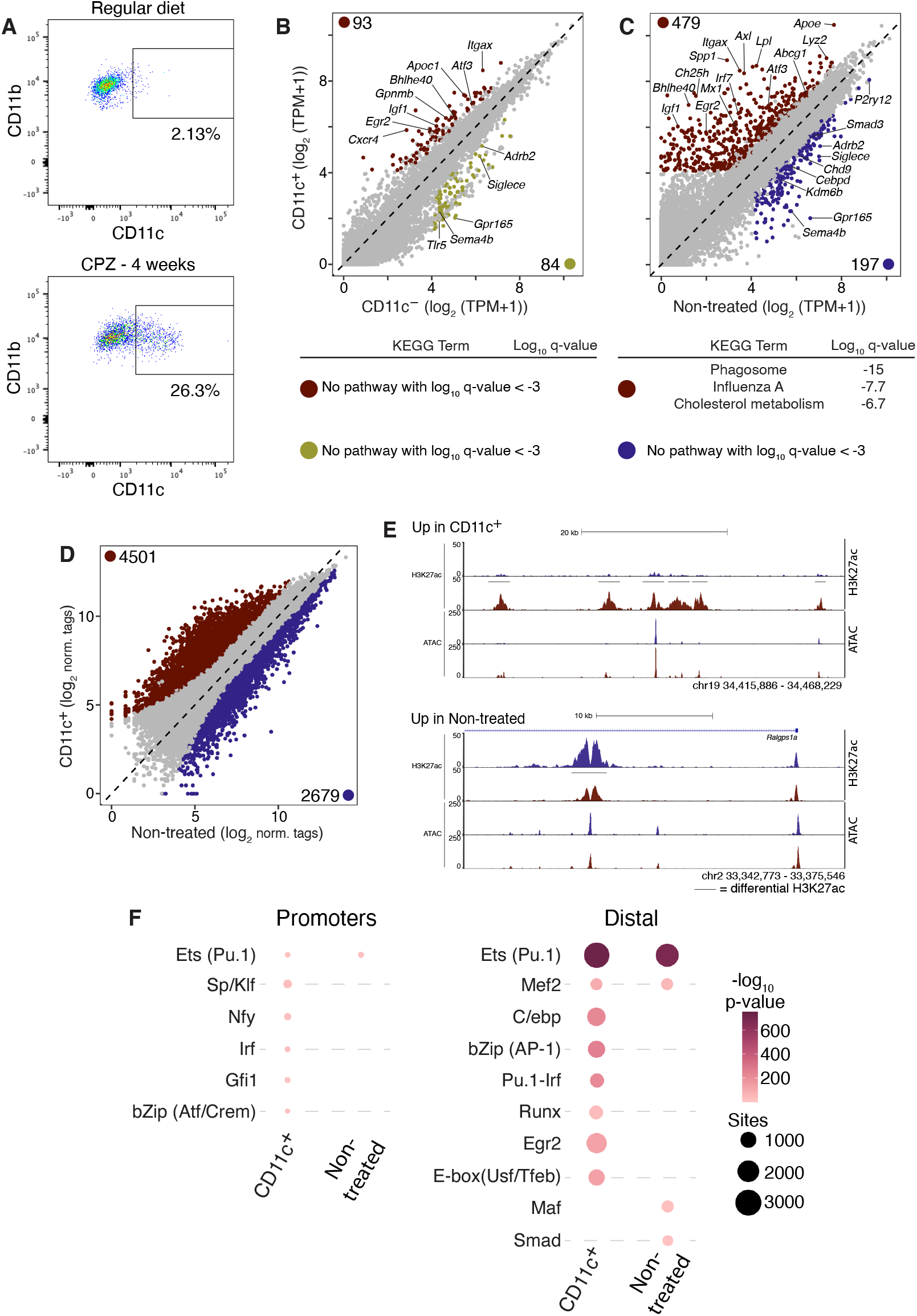
Transcriptional and epigenomic features of CD11c^+^ microglia phenotype associated with chronic myelin lesions. **(A)** Flow cytometry data depicting detection of CD11c^**+**^ and CD11c^**−**^ microglia in the brain of mice fed cuprizone (CPZ) diet for 4 weeks. **(B, C)** Scatterplots of RNA-seq data comparing CPZ CD11c^**+**^ and CPZ CD11c^**−**^ microglia (B) and CPZ CD11c^**+**^ and non-treated/CD11c^**−**^ (C) microglia isolated from healthy mice. Differentially expressed genes identified by DESeq2 (log2 FC ≥ 0.58, FDR ≤ 0.01) are colored coded. Associated KEGG pathways analysis results are also appended. **(D)** Scatterplot of H3K27ac abundance at distal CREs, comparing CPZ CD11c^**+**^ and non-treated/CD11c^**−**^, as assessed by ChIP-seq. Differentially enriched CREs identified by DESeq2 (FDR-adjusted p-value ≤ 0.05, two-fold or greater tag difference) are colored coded. **(E)** UCSC browser displays of H3K27ac and ATAC-seq data at representative loci of genomic elements that exhibit significantly higher (top) or lower (bottom) levels of H3K27ac in CPZ CD11c^**+**^ vs non-treated/CD11c^**−**^ microglia. **(F)** Table summarizing results from de novo DNA motifs analyses performed on differentially active promoters and distal CREs identified in C and D, respectively.

The high prevalence of Egr2 and E-box motifs at upregulated CREs in CPZ CD11c^**+**^ microglia were not prominent features associated with the motif signatures associated with the LPS model. This further supports the general conclusion that microglial gene regulation is inherently linked to the surrounding state of the brain microenvironment. Yet, dominant antiviral gene signatures and enrichment of Pu.1 and AP-1 and C/ebp motifs at CREs detected in both CPZ CD11^+^ and LPS-associated microglia suggests that some transcriptional regulators may be commonly activated across different types of disturbances afflicting the brain. Thus, we next sought to dissociate probable genes and CREs whose regulation occurs as part of a broad, shared program of inflammatory activity from those that may be more susceptible to context-dependent regulation. Overall transcriptome comparisons of the different LPS and CPZ subsets isolated supported similarities and differences between the CPZ and LPS subsets (Fig. 6A). Specifically, the CPZ and LPS 48h microglia are most similar to one another, both clustering apart from the LPS early timepoints (i.e., 3, 6 and 24h). We next performed more systematic comparisons of the CPZ and LPS subsets to dissociate potential genes subject to pan-regulation from those whose expression is more perturbation-specific. For this, we overlapped genes that are significantly induced in at least one of the LPS subsets compared to microglia from the healthy brain with those induced in CPZ-CD11c^**+**^ microglia *vs* basal state (log2 FC ≥ 0.58, FDR-adjusted p-value ≤ 0.01). Overall, 263 genes were commonly induced in both models, and these included several genes involved in antiviral immunity (Fig. 6B to D; Table S4). In contrast, LPS-biased genes included 1,293 genes and these were frequently linked with TNF/Nf-κB signaling and regulation of the cell cycle process. Lastly, CPZ CD11c^**+**^ microglia upregulated 216 genes in a relatively specific manner. Coherent with observations made earlier, these included several genes coding for effectors of cholesterol metabolism, including *Apoe* and *Lpl*.

**Figure 6.**
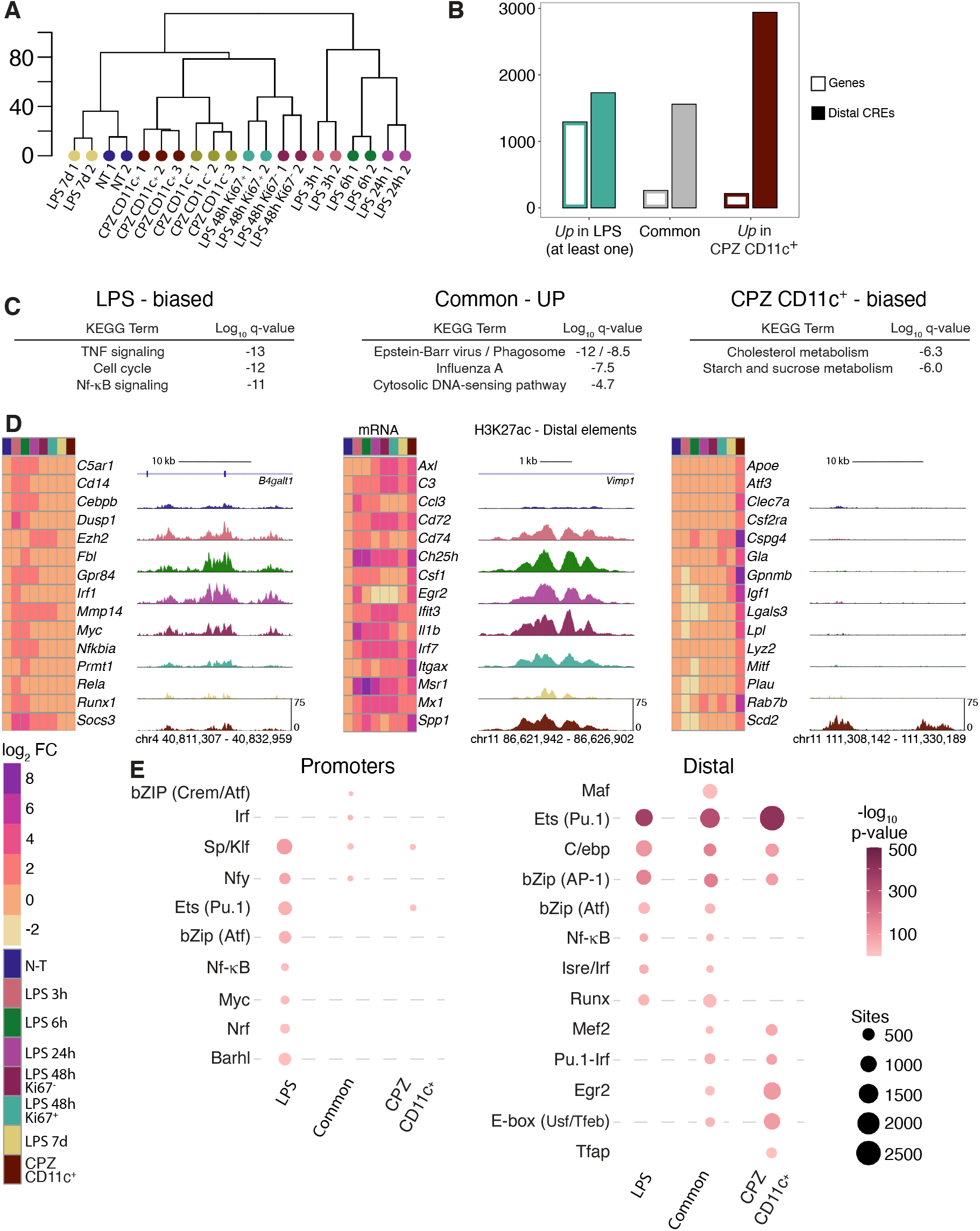
Activation of context-common and context-specific genomic regulatory elements associated with distinct inflammatory microglial phenotypes. **(A)** Dendrograms depicting overall similarities of gene signatures of different LPS and CPZ-associated inflammatory microglial subsets. **(B)** Bar graphs representing number of genes (empty bars) and distal CREs (solid bars) that are preferably expressed in at least one LPS subset vs CPZ CD11c^+^ microglia, and vice versa. **(C)** KEGG pathways analyses for genes identified in B. **(D)** Heatmaps depicting mean values of mRNA log2 FC for microglial inflammatory phenotypes and UCSC browser displays of H3K27ac data of genomic loci, all identified in B. **(E)** Table summarizing results from de novo DNA motifs analyses performed on differentially active promoters and distal CREs identified in B.

Given the evidence for perturbation-biased gene expression, we next interrogated the extent to which promoter-distal regulatory elements activated in inflammatory microglia could also segregate according to pathological context. Overlapping analyses revealed that 1559 elements commonly gained H3K27ac in at least one LPS condition and CPZ-CD11c^**+**^ microglia, and these were highly enriched for Pu.1, bZip, and C/epb motifs (Fig. 6B, D, E). Also, while 1731 elements were higher for H3K27ac in at least one LPS condition (i.e., LPS-biased), 2942 were preferably activated in CPZ CD11c^**+**^ microglia (Fig. 6B, D). Notably, DNA motif analyses suggested contribution stronger input from Nf-κB and Runx for the LPS-biased response, and stronger contribution from Egr2, Mef2 family members and Tfeb/Usf factors for the CPZ CD11c^**+**^-biased activity. Therefore, these data provide evidence that microglia in different pathological brain conditions leverage different configurations of signaling and transcriptional effectors, which ultimately contribute to perturbation-specific profiles of gene expression.

### Mef2a activity promotes the microglial CD11c^+^ phenotype

To assess the functional relevance of the TFs implicated by the motifs analyses above, we interrogated the role of Mef2a in the regulation of CD11c^**+**^ microglia differentiation and/or function in the CPZ model. Observations supporting this selection includes 1) the high enrichment for Mef2 motifs at CPZ CD11c^**+**^-biased CREs, 2) the high expression of *Mef2a* mRNA in CPZ CD11c^**+**^ microglia (Fig. 7A), and 3) Mef2 factors have previously been implicated in the regulation of phagocytic activity of macrophages in vitro(36). Injection of Tamoxifen in male and female Cx3cr1^CreERT2/WT^::Mef2a^fl/fl^ mice led to robust inhibition of *Mef2a* mRNA expression (Fig. 7B). Deletion of *Mef2a* in microglia did not impact Iba1 microglial density in cortical or striatal regions in the healthy brain (Fig. 7C, D). However, absence of *Mef2a* gene expression led to a 35-40% decrease in the proportion of CD11c^**+**^ microglia in the brain of mice fed CPZ for 4 weeks (Fig. 7E, F); expression levels were also lower in Cx3cr1^CreERT2/WT^::Mef2a^fl/fl^ mice (Fig. 7E, F). Of note, abrogating Mef2a expression did not impact the proportion of proliferating Ki67^**+**^ microglia, suggesting that not all aspects of microglial cell activity are disrupted in absence of this transcription factor. Overall, these observations suggest that Mef2a activity promotes differentiation and/or sustains the CD11c^**+**^ microglial subset associated with chronic myelin lesions in the brain.

**Figure 7.**
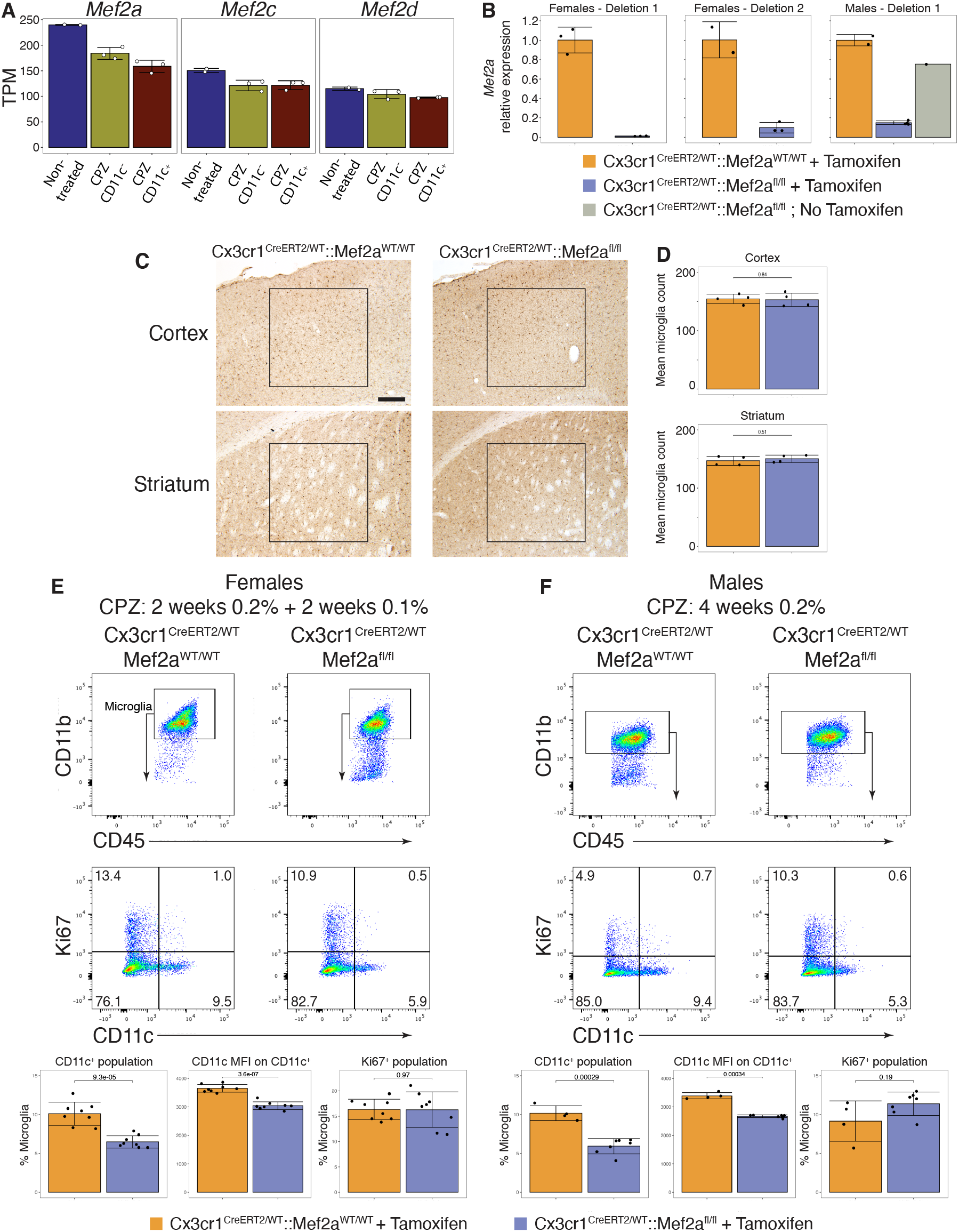
Mef2a promotes CD11c expression in microglia during demyelinating brain lesions. **(A)** mRNA levels for *Mef2* family members expressed in non-treated and CPZ-associated microglia (Transcripts Per Million (TPM) normalization). **(B)** qPCR data depicting *Mef2a* mRNA expression, normalized to *36b4* mRNA levels, in microglia isolated from Cx3cr1^CreERT2/WT^::Mef2a^WT/WT^ and Cx3cr1^CreERT2/WT^::Mef2a^fl/fl^ mice 8 weeks after Tamoxifen injection. **(C)** Representative microphotographs depicting Iba1 immuno-labeling in cortical and striatal areas of healthy male Cx3cr1^CreERT2/WT^::Mef2a^WT/WT^ and Cx3cr1^CreERT2/WT^::Mef2a^fl/fl^ mice. Scale bar: 200 μM **(D)** Quantification of data presented in C. **(E)** Representative flow cytometry data depicting expression of CD11c and Ki67 in microglia of mice fed CPZ diet for 4 weeks. **(F)** Quantification of data presented in D. MFI: Median Fluorescence Intensity.

## Discussion

Here we report on a comprehensive characterization of epigenomic and transcriptional mechanisms associated with microglial inflammatory activity in acute and chronic pathological brain conditions. Our work suggests that axes of signaling pathways – transcription factors interact with sets of common but also context-specific genomic regulatory elements to couple the transcriptional output of microglia to the surrounding perturbed brain microenvironment. Computational analyses of these regulatory elements also identified combinations of transcriptional regulators that may underlie context-dependent microglial inflammatory responses. These arrangements likely operate through a two-tier system. First, Pu.1, AP-1 and C/epb family members appear to provide dominant, pan regulatory input, independently of neuroinflammatory settings. Their activity then combines with more restricted, context-biased signals, such as Nf-κB, Irf and Runx to coordinate anti-microbial, alarming responses, or Egr2, Mef2 and E-box TFs for tissue-remodeling functions. Overall, these studies extend the role of genomic regulatory elements in coordinating context-specific transcriptional responses to inflammatory microglia in the brain. They also provide novel insights into the molecular mechanisms that coordinate the ability of these cells to promptly adapt their functions upon significant breakdowns in brain homeostasis.

Time-course sampling of microglia using the model of systemic LPS injection is a robust strategy to monitor the dynamic transcriptional mechanisms of gene expression triggered as microglia mobilize to engage in inflammatory activity. Overall, our results confirm that microglia rapidly detect signaling cues linked to a systemic inflammatory response and promptly proceed with a first wave of transcription of inflammatory genes within 3 hours(1, 23). Furthermore, while the number of genes induced as part of the first wave is substantial, the number of promoter-distal elements that gained H3K27ac was comparatively low. DNA motifs analysis also revealed rather weak, sparse enrichment for Nf-κB, Irf, and bZip-binding factors like AP-1 and Atf family members at these sites. Together, these observations suggest that the first wave may be part of a general transcriptional alarm response that primarily relies on the direct control of the transcriptional machinery at gene promoters, perhaps through pause-release-targeting mechanisms. They also further highlight that microglia, as innate immune cells, are inherently programmed to rapidly trigger and coordinate inflammatory responses in the brain. Shortly after the first wave follows the transcriptional induction of effectors of protein synthesis. At this point, the underlying mechanism of transcriptional regulation becomes more complex, as evidenced by the marked increase of promoter-distal elements that displayed elevated H3K27ac at 6 hours. This enhanced response likely results from sustained, albeit also probably stronger, input from Nf-κB, AP-1, Atf, and Irf family members and recruitment of additional input. These may include C/epb family members, Runx, and E-box-binding factor that act specifically at promoter-distal elements, and Myc acting preferentially at promoters. Subsequently, by the end of the first day, a large subset of microglia deploys a cell cycle gene program, leading to a peak of inflammatory microglia cell proliferation between 24 and 72 hours. Finally, the neuroinflammatory response is largely resolved by the seventh day, with the microglial gene signature and H3K27ac genome-wide profiles returning to pre-insult states. This is likely due to the absence of sustained inflammatory input inherent to the transient nature of the LPS model used. It is also possible that resolving mechanisms, such as Il-10 signaling, actively contribute to the resolution of microglial inflammatory activity in the systemic LPS model(25).

Our results highlight that several signaling pathways implicated in microglial homeostatic activity become downregulated during an inflammatory response. For example, we recovered evidence that mRNA expression of *Maf* decreases in the LPS-associated response, coherent with previous studies on inflammatory microglia gene regulation(9). However, we also provide novel evidence indicating that their functional transcriptional activity is also downregulated as revealed by enrichment of their motifs at distal CREs that lose H3K27ac during the pre-proliferative inflammatory stage. We note that the Smad motifs was also frequently significantly enriched at such elements, pointing to an inhibition Tgf-β1 signaling in inflammatory microglia. This latter observation is further supported by RNA-seq data that showed a decreased in *Sall1* mRNA, that latter of which is a key transcriptional co-repressor whose expression in microglia is positively regulated by Tgf-β1 cytokine(22, 37).

These studies provide further support for an intricate relationship between inflammation, transcription and microglial proliferation(31, 32). To this end, expression levels of several fundamental signaling regulators of macrophages proliferation are dynamically regulated by the transcriptional process in the LPS model. This occurs within the first 24 hours of the LPS-associated response, well preceding the peak of microglia population expansion. For example, as part of the first wave, microglia upregulate mRNAs coding for Csf1, a potent growth factor for microglia(38), and Runx1, a transcription factor involved in microglial and macrophage cell cycling(39, 40). As Runx1 is an effector of downstream of Csf1r(40), these observations suggest that Csf1 may act on microglia in autocrine fashion, stimulating high Runx1 activity to facilitate proliferation. We also note that effective activity of Myc and E2f1, which are key transcriptional regulators of cell proliferation(41, 42), is also likely calibrated by their expression at the transcriptional level during the first and third wave, respectively. In addition, the rapid decrease in the expression of *Maf* mRNA agrees with work in primary macrophages that showed that joint inhibition of Maf transcription factors enhances the proliferative capabilities of these cells(43, 44). Therefore, transcriptional modulation of the expression of pro-and inhibitory regulators likely leads to a situation where activation thresholds of microglial proliferation are reached and exceeded, thus enabling the massive proliferation of the microglial population.

In contrast to the acute LPS model, microglia in the demyelinating brain engage in sustained, heightened immune activity. Also unlike the LPS-based activity, which is dominated by strong induction of antiviral and Tnf-associated gene programs, CPZ-CD11c^**+**^ microglia display more prominent expression of genes linked to phagosomal cell biology. Altogether, this is highly coherent with the potent capabilities for debris elimination associated with CD11c^**+**^ microglia in chronic brain lesions(24). Notably, analyses of CREs suggested that Egr2, Mef2 members, and E-box binding factors such as Tfeb/Usf family members may provide key regulatory input required for the CD11c^**+**^ polarization. Multiple lines of evidence suggest functional roles for these TFs in such context. First, abrogating Mef2a expression decreased the proportion of CD11c^**+**^ microglia but had no apparent effects of cell proliferation. Second, Tfeb family members, including Tfeb, Tfe3, and Mitf, which are all reliably expressed in CPZ-CD11c^**+**^ microglia (Table S5), are important positive regulator of lysosomal biogenesis(45, 46); evidence also suggests that Usf factors may promote this function as well(47). Lastly, high Egr2 factor activity licenses the so-called macrophage alternative phenotype(48, 49), which also possesses potent tissue remodeling properties. Upstream regulators of these factors in microglia remain to be precisely defined, however. For example, elevated Egr2 expression in alternatively polarized macrophages derives from high Stat6 transcriptional activity, but the Stat6 motif was never prevalent in any of the CPZ microglia subsets interrogated here(48, 49). Nevertheless, we can speculate based on literature on the identity of distinct signals that may promote differentiation of the CD11c^**+**^ subset during lesions. For example, Trem2 signaling promotes both expression of *Itgax* mRNA, the gene coding for CD11c and microglia cell population expansion in the CPZ model(50). Also, we note that while Igf-1 promotes microglial proliferation after spinal cord injury(51), augmented expression of *Igf1* mRNA in CPZ-CD11c^**+**^ microglia may also perhaps act in autocrine fashion to promote additional cell functions for the CD11c^**+**^ subset.

Multiple outstanding questions emerge from this work. For example, our results implicate broad activity from large families of transcription factors, including Nf-κB, AP-1, Atf, Irf, Tfeb and Mef2 to name a few. However, the specific contribution of their different family members remains to be defined. In line with this, the extent to which their effective input changes quantitatively and/or qualitatively over time as microglia proceed through the response is not known. For example, we observed *de novo* upregulation of *Irf7* mRNA in microglia in the LPS model; it is thus unlikely that this factor participates to the original Irf-driven response, but it may be important at later timepoints to sustain the Irf-mediated gene transcription. Finally, the extent to which Egr2, Mef2, and E-box binding factors also regulate the CD11c^**+**^ microglial phenotype in other lesion contexts, including neurodegenerative disorders such as AD, remains to be investigated. Whether their activity can be manipulated to achieve therapeutic benefits with neurodegenerative disorders remains to be tested as well.

## Material and Methods

### Mice

C57BL/6J (The Jackson Laboratories, stock 00664), Mki67^tm1.1Cle^ (Ki67-RFP; The Jackson Laboratories, stock 029802) and Cx3cr1^tm2.1(cre/ERT2)Jung/J^ (Cx3cr1^CreERT2/CreERT2^; The Jackson Laboratories, stock 020940) were originally purchased from The Jackson Laboratory. Mef2a^fl/fl^ mice(52) were generously and kindly donated by Dr Eric Olson from UT SouthWestern (Dallas, Texas, USA) and bred with Cx3cr1^CreERT2/CreERT2^ to create the Cx3cr1^CreERT2/WT^::Mef2a^fl/fl^ mouse line. Internal colonies for each of these mouse lines were established upon reception of breeding pairs at the animal research facility of Centre de Recherche du Centre Hospitalier Universitaire de Québec-Université Laval. Unless otherwise specified, males were used throughout this study. All adult mice were of 2 to 4.5 months of age at sacrifice. Mice had ab libitum access to food and water. All animal procedures were approved by Université Laval’s animal care and ethics committee (CPAUL3) and performed in compliance with ethical regulations and guidelines of the Canadian Council on Animal Care.

### Mouse genotyping

For genotyping, DNA was extracted from ear punches using the HotSHOT method. For the Mef2a^fl/fl^ construct, PCR reaction of 15 μl was setup using the following conditions: 1 μl of genomic DNA, 0.6 μM of each primer, 200 μM of dNTP, 1X standard buffer and 1.5 units of HotStart Taq (NEB, M0495). Primer sequences were described previously(52): Gtm2ACKO-SA-FWD, 5’-GGTAGCTCAGGTGTCACTTCTTG-3’; Gtm2ACKO-KO-REV, 5’-CACTTTACATCCCAATAGCAGCC-3’; Gtm2ACKO-LA-REV, 5’-CTCATCCATTTATGGCTGTGTC-3’. Expected PCR product size for WT band is 267 base pairs (bp), loxP band 367 bp and KO 500 bp. Genotyping of the Cx3cr1^CreERT2^ construct used DNA extraction and PCR conditions similar to those used for *Mef2a* gene. PCR primers were obtained from The Jackson Laboratory: WT FWD 5-AGCTCACGACTG CCTTCTTC-3’, common 5’-ACGCCCAGACTAATGGTGAC-3’ and mutant FWD 5’-GTTAATGACCTGCAGCCAAG-3’. Expected PCR product sizes for WT band is 151 bp and mutant band 230 bp.

### LPS treatment and cuprizone diet

LPS (1 mg/kg; from Escherichia coli, serotype 055:B5; Sigma-Aldrich, L-2880), diluted in a volume of 100 μl of 0.9% saline solution was administered intraperitoneally. For all except one experiment with cuprizone diet, cuprizone (Sigma-Aldrich, 14690) was mixed with regular ground chow (0.2% wt/wt) and fed ab libitum to mice for 4 weeks(53). For one experiment (i.e., Fig.7E, females), mice were initially fed cuprizone at 0.2% for 2 weeks and then 0.1% for an additional 2 weeks. In all cases diet was changed every 2 days.

### Tamoxifen-induced *Mef2a* deletion in microglia

To delete *Mef2a* gene expression in microglia, we administered Cx3cr1^CreERT2/WT^:: Mef2a^fl/fl^ mice 4 injections of Tamoxifen (Sigma-Aldrich, T5648) over 4 consecutive days (1 injection/day), starting on postnatal day 36. Tamoxifen was dissolved in 10% EtOH, 90% corn oil (Sigma-Aldrich, C8267) and injected at a final dose of 100 mg/kg, in a volume of 100 μl. Control mice consisted of age- and sex-matched Cx3cr1^CreERT2/WT^:: Mef2a^WT/WT^ that underwent to same Tamoxifen administration regimen. Mice were then left undisturbed for at least 6 weeks after the last injection. To validate deletion efficiency, microglia were first isolated as described below, RNA then isolated (Direct-zol RNA Micro Prep, Zymo Research), and cDNA generated (OneScript Hot cDNA Synthesis Kit, ABM) based on manufacturer’s instructions. *Mef2a* mRNA expression was then assessed by qPCR (PowerTrack SYBR Green Master Mix, Applied Biosystems, LightCycler 480 Instrument II, Roche), along with that of *36b4* mRNA for data normalization, using the delta delta Ct calculation method. Primers for *Mef2a* mRNA were FWD 5’-AGACAAGGTGACTGAAAATGGG-3’ and REV 5’-CAGAGCACACTGAGTTCATAGG-3’, and for *36b4* mRNA FWD 5’-GCTCGACATCACAGAGCAGG-3’ and REV 5’-CCGAGGCAACAGTTGGGTAC-3’.

### Microglia isolation and flow cytometry analysis

Mice were first euthanized with an overdose of ketamine-xylazine administered by intra-peritoneal injection and subsequently perfused transcardially with PBS. Brain were then removed from the skull and immediately homogenized in a buffer solution (HBSS 1X (Life Technologies, 14175-095), 1% BSA (Sigma-Aldrich, A9647), 1mM EDTA (Invitrogen, 15575020), 1 mM sodium butyrate (Sigma-Aldrich, B5887), 1 μM Flavoperidol (Sigma-Aldrich, F3055)) by gentle mechanical dissociation on ice, using a 7 ml dounce tissue grinder (DWK Life Sciences, 357542) as performed in Gosselin et al. 2014(54). Microglia were then enriched by Percoll (Cytivia, 17089101) gradient, washed and labeled with LIVE/DEAD Fixable Yellow Dead Cell Stain Kit (Invitrogen), then incubated with a CD16/CD32 receptor blocking antibody (1:50, BioLegend, 101302, clone 93) for 10 min, and then incubated 1:100 with antibodies directed against cell surface proteins CD11b-BV421 (BioLegend, 101251, clone M1/70), CD45-Percp-Cy5.5 (BioLegend, 103132, clone 30-F11), and CD44-APC-Cy7 (BioLegend, 103028, clone IM7) and CD11c (BioLegend, 117310, clone N418) for 30 min. For Ki67 assessment, microglia were then further fixed with 1% PFA for 10 min at RT, permeabilized with eBioscience Foxp3 / Transcription Factor Fixation/Permeabilization Concentrate and Diluent (Invitrogen) and stained with anti-Ki67-Alexa 488 antibody (Invitrogen, 53-5698-82, clone SolA15). Microglia were either analyzed with BD LSR cell analyzer or sorted in staining buffer with BD Aria II cell sorter (100 μm nozzle). Flow cytometry results were analyzed using FlowJo software (BD).

### RNA-seq library preparation: Isolation and fragmentation of poly(A)

For RNA-seq, we generated at least 2 independent biological replicates per conditions (n = at least 2). Following sorting, 200,000 microglia were pelleted and put into 100 μl lysis/Oligo d(T)25 Magnetic Beads binding buffer (100 mM Tris-HCl ph 7.5, 500 mM LiCl, 10 mM EDTA pH 8.0, 1% LiDS, 5 mM DTT, in water) and stored at -80 °C until processing. For human brain cortical RNA, 500 ng of RNA was diluted with proper volume of 2x lysis/ Oligo d(T)25 Magnetic Beads binding buffer to a final concentration of 1x lysis/Oligo d(T) Magnetic Beads binding buffer. For sequencing library preparation, samples were incubated with 20 μl Oligo d(T)25 Magnetic Beads (NEB, S1419S) in PCR strips for selection of polyadenylated (poly(A)) mRNA as follows: 2 min at 65 °C on a PCR cycler and then 10 min at room temperature (RT). Samples were then placed on a collection magnet at RT and washed serially with 180 μl of washing buffer 1 (RNA-WB1; 10 mM Tris-HCl pH 7.5, 0.15 M LiCl, 1 mM EDTA pH 8.0, 0.1% LiDS, 0.1% Triton X-100, in water) and then washing buffer 3 (RNA-WB3; 10 mM Tris-HCl, 0.15 M NaCl, 1 mM EDTA pH 8.0, in water). RNA was eluted from Oligo d(T)25 beads by resuspending the beads in 50 μl of elution buffer (EB; 10 mM tris-HCl pH 7.5, 1 mM EDTA pH 8.0, in water) followed by incubation at 80 °C for 2 min on a PCR cycler. The samples were then placed back on the magnet, and the elution buffer supernatant containing the poly(A) RNA was carefully collected and placed on ice.

A second round of poly(A) RNA selection was then performed. Eluted Oligo d(T)25 beads were first washed on a magnet with 150 μl EB and then 150 μl 2X Oligo d(T)25 binding buffer (2x DTBB; 20 mM Tris-HCl pH 7.5, 1 M LiCl, 2 mM EDTA pH 8.0, 1% LiDS, 0.1% Triton X-100) and resuspended in 50 μl of 2x DTBB. Resuspended Oligo d(T)25 beads were then added to isolated RNA and incubated at 65 °C and at RT as described above. Beads were then placed on a magnet and washed once with 150 μl of RNA-WB1 and then with 30 μl of 1x First-Strand buffer (made from 5x stock; Life Technologies 18080-044) and transferred in a new PCR strip. Beads were collected on a magnet and resuspended in 10 μl of 2x SuperScriptIII buffer (Invitrogen, 1x First-Strand buffer, 10 mM DTT in water). Resuspended beads were then incubated at 94 °C for 9 min to fragment poly(A) RNA. Beads were then placed back on a magnet, and the supernatant containing poly(A) RNA was carefully collected and placed on ice.

### RNA-seq library preparation: Synthesis of cDNA

RNA in 2X SuperScript III buffer was incubated for 1 min at 50 °C on a PCR cycler with 2.5 μl of the following mix: 1.5 μg Random Primer (Invitrogen, 48190-011), 10 μM Oligo d(T)20 primer (Invitrogen, 18418020), 10 units SUPERase•In RNase Inhibitor (Invitrogen), 4 mM dNTP mix (Invitrogen, 18427088), in water. Samples were then immediately placed on ice for 5 min. First-strand synthesis was then performed by incubation at 25 °C for 10 min and 50 °C for 50 min on a PCR cycler with 7.6 μl of the following mix: 0.2 μg actinomycin D (Sigma-Aldrich, A1410), 13.15 mM DTT (Thermo Scientific), 0.026% Tween-20, 100 units SuperScript III Reverse Transcriptase (Invitrogen, 18080-044), in water. After incubation, RNA/DNA complexes were isolated by adding 36 μl of RNAClean XP beads (Beckman Coulter, A63987) and incubated for 10 min at RT and then 10 min on ice. Samples were then placed on a magnet and beads were washed twice with 150 μl of 75% EtOH. Following washings, beads were air-dried for 10-12 min and eluted with 10 μl of water.

Second-strand synthesis was then performed. RNA/DNA samples, in 10 μl of water, were incubated for 2.5 hours at 16 °C with 5 μl of the following mix: 3x Blue Buffer (Enzymatics, B0110), 1.0 μl PCR mix (Affymetrix, 77330), 2.0 mM dUTP (Affymetrix, 77206), 1 unit RNAse H (Enzymatics, Y9220L), 10 units DNA Polymerase I (Enzymatics, P7050L), 0.03% Tween-20, in water). DNA was then purified by addition of 1.5 μl Sera-Mag SpeedBeads Carboxyl Magnetic Beads (Cytivia, 45152105050250), resuspended in 30 μl 20% PEG 8000/2.5 M NaCl, incubated at RT for 15 min and placed on a magnet for two rounds of bead washing with 80% EtOH. Beads were then air dried for 10-12 min and DNA was eluted from the beads by adding 40 μl of water. Supernatant was then collected on a magnet and placed on ice or stored at -20 °C until DNA blunting, poly(A)-tailing, and adapter ligation (see below).

### Chromatin immunoprecipitation

Chromatin immunoprecipitation for histone modification H3K4me3 and H3K27ac was performed as follows, using ∼500,000 microglia per assay and n=2 independent biological replicates per conditions. First, microglia were briefly thawed on ice and lysed by incubation in 1 ml of lysis buffer (0.5% IGEPAL CA-630, 10 mM HEPES pH 7.9, 85 mM KCl, 1 mM EDTA pH 8.0, in water) for 10 min on ice. Lysates were centrifuged at 800 RCF for 5 min at 4 °C, and pellets were resuspended in 200 μl of sonication/ immunoprecipitation buffer (10 mM Tris-HCl pH 7.5, 100 mM NaCl, 0.5 mM EGTA, 0.1% Deoxcycholate, 0.5% Sarkosyl, in water). Sonication was performed with a Bioruptor Standard Sonicator (Diagenode) and consisted of two rounds of 15 min each, alternating stages of 30 sec “sonication-on” with 60 sec “sonication-off”. Twenty-two microliters of 10% Triton-X were then added to samples (1% final concentration) on ice, and lysates were cleared by centrifugation for 5 min at 18,000 RCF at 4 °C. Two microliters of supernatant were then set aside for input library sequencing controls. Supernatants were then immunoprecipitated on a rotator for two hours at 4 °C with antibody pre-bound to 17 μl Protein A Dynabeads (Invitrogen, 10001D). The following antibodies was used: H3K27ac (Active Motif, 39685), 2.5 μg per sample.

Immunoprecipitates were washed three times each on ice with ice-cold wash buffer I (150 mM NaCl, 1% Triton X-100, 0.1% SDS, 2 mM EDTA pH 8.0 in water), wash buffer III (10 mM Tris-HCl, 250 mM LiCl, 1% IGEPAL CA-630, 0.7% Deoxycholate, 1mM EDTA in water) and TET (10 mM Tris-HCl pH 7.5, 1 mM EDTA pH 8.0, 0.1% Tween-20, in water) and eluted with 1% SDS/TE at RT in a final volume of 100 μl.

Reverse-crosslinking was then performed on immunoprecipitates and saved input aliquots. First, 6.38 μl of 5 M NaCl (final concentration 300 mM) was added to each sample immersed in 100 μl SDS/TE. Crosslinking was then reversed by overnight incubation at 65 °C in a hot air oven. Potentially contaminated RNA was then digested for one hour at 37 °C with 0.33 mg/ml RNase A, proteins were digested for one hour at 55 °C with 0.5 mg/ml proteinase K, and DNA was extracted using Sera-Mag SpeedBeads.

### RNA-seq and ChIP-seq final library preparation

Sequencing libraries were prepared from recovered DNA (ChIP) or generated cDNA (RNA) by blunting, A-tailing, and adapter ligation as previously described using barcoded adapters (NextFlex, Bioo Scientific)(22, 54). Prior to final PCR amplification, RNA-seq libraries were digested by 30 min of incubation at 37 °C with Uracil DNA Glycosylase (final concentration of 0.134 units per μl of library volume; UDG, Enzymatics, G5010L) to generate strand-specific libraries. Libraries were PCR-amplified for 12-15 cycles and size selected for fragments (200-400 bp for ChIP-seq, 200-500 for RNA-seq) by gel extraction (10% TBE gels, Invitrogen, EC62752BOX). RNA-seq and ChIP-seq libraries were single-end sequenced for 76 cycles on an Illumina HiSeq 4000 (Illumina, San Diego, CA) according to manufacturer’s instruction.

### Assay for Transposase-Accessible Chromatin-sequencing (ATAC-seq)

100,000 isolated microglia were lysed in 50 μl lysis buffer (10 mM Tris-HCl ph 7.5, 10 mM NaCl, 3 mM MgCl2, 0.1% IGEPAL, CA-630, in water) on ice and nuclei were pelleted by centrifugation at 500 RCF for 10 min. Nuclei were then resuspended in 50 μl transposase reaction mix (1x Tagment DNA buffer (Illumina, 15027866), 2.5 μl Tagment DNA enzyme I (Illumina, 15027865), in water) and incubated at 37 °C for 30 min on a PCR cycler. DNA was then purified with Zymo ChIP DNA concentrator columns (Zymo Research, D5205) and eluted with 10 μl of elution buffer. DNA was then amplified with PCR mix (1.25 μM Nextera primer 1, 1.25 μM Nextera primer 2-bar code, 0.6x SYBR Green I (Invitrogen, S7563), 1x NEBNext High-Fidelity 2x PCR MasterMix, (NEBM0541) for 9 cycles, run on a gel for size selection of fragments (160-290 bp), extracted from the gel and single-end sequenced for 76 cycles on a HiSeq 4000.

### Postnatal microglia and PLX5622 microglia datasets

Data from postnatal microglia and PLX5622-associated microglia were generated as part of a previous study (Belhocine et al., 2022(31); Gene Expression Omnibus platform series GSE166236).

### Sequencing data analysis

#### Preprocessing

FASTQ files from sequencing experiments were mapped to the mouse mm10 reference genome. STAR with default parameters was used to map RNA-seq experiments(55). Bowtie2 with default parameters was used to map ChIP-seq and ATAC-seq experiments(56). HOMER was used to convert aligned reads into “tag directories” for further analyses(26).

#### RNA-seq

For differential expression analyses, a table read count including relevant samples was first created using the *analyzeRepeats*.*pl* program of HOMER (v4.11; http://homer.ucsd.edu/homer/) with the following parameters: *rna -noadj - condenseGenes -count exons -pc 3*. The resulting dataset was used as input file for DESeq2. Wald Test was performed with DESeq2 to identify differentially expressed genes between groups of interest, using the following parameters: lfcThreshold = 0.58, alpha = 0.01. For final data plotting and interpretation purposes and KEGG pathway analyses (see below), we removed non-coding RNAs and pseudogenes and retained only protein-coding mRNAs that have a Transcript Per Million (TPM)-normalized value of 16 or greater. The base-2 logarithm of the TPM values was taken after adding a pseudocount of 1 TPM to each gene. TPM datasets were generated using HOMER’s *analyzeRepeats*.*pl* program with the following parameters: *rna -tpm -condenseGenes - count exons*. Global raw and TPM-normalized RNA-seq data for all samples used in this study are provided in Table S5.

#### H3K27ac ChIP-seq regions calling

For each condition, the repertoire of genomic regions positive for presence of H3K27ac were first generated using HOMER’s *getDiffPeakReplicate*.*pl* program, using all replicates and related input of a condition of interest. Parameters specified for H3K27ac were as follow: *-genome mm10 -region - size 500 -min Dist 1000 -L1*. For differential enrichment assessments, relevant regions files were then first merged together with HOMER’s *mergePeaks*.*pl* program with *-size given* parameter, and tag counts of each replicate annotated to each region using HOMER’s *annotatePeak*.*pl* program with *mm10 -raw* parameters used. For H3K27ac, data matrix was then used as input for HOMER’s *getDiffExpression*.*pl* program, which leveraged R/DESeq2 to assess differential enrichment. Regions with two-fold differences or greater in normalized tag counts at a false-discovery rate (FDR)-adjusted p-value of 0.05 or lower were considered as differentially enriched between conditions compared. For data interpretation and plotting, only regions with 16 or more normalized reads were considered. For read count tables output, each experiment was normalized to 10 million total reads (*-simpleNorm* default parameter specified in *getDiffExpression*.*pl*).

#### ATAC-seq accessible chromatin region identification

Tag directories of condition replicates were first merged using HOMER’s *makeTagdirectory*.*pl*. Regions of accessible chromatin were then called using HOMER’s *findPeaks*.*pl* program with the *- style factor* parameter.

### De novo motif enrichment analyses

#### Promoters analyses

For promoter analyses, de novo motif enrichment analysis was performed on the area of open chromatin defined by ATAC-seq (i.e., ATAC-seq “peaks”) located within -1000 to +500 bp genomic regions that encompass a transcriptional start sites (TSS) of protein-coding genes of interest (mm10, RefSeq annotation). For this, tables of genomic coordinates of promoters were first generated with HOMER’s *annotatePeaks*.*pl* program in *tss* mode, using as input the list of NM RefSeq gene identifiers of relevant subgroups of genes (see Results), and specifying the following parameters: *-list <relevant NM list*.*txt> -size -1000,500*. Output was then curated to retain only gene regions included in the original NM list .txt file. Following this, relevant regions for motifs analysis were recovered using the *intersect* program of BEDTools(57) against ATAC-seq peak files. Finally, that latter BEDTools output file was used as input for HOMER’s *findMotifsGenome*.*pl* command with the following parameters: *-size 300 -mask*. Background sequences for differential enrichment assessment were generated randomly by the program. Finally, motifs that were present in at least 5% of intersected ATAC-seq “peaks” and that had a score of at least 0.8 were retained for data interpretation.

#### Promoter-distal regulatory elements analyses

Promoter-distal CREs were first defined as H3K27ac-positive regions located outside of the 1000 to +500 bp genomic regions that encompass any TSS of protein-coding genes (mm10, RefSeq annotation). DNA motif enrichment analyses for promoter-distal CRE was centered ATAC-seq “peaks” located within CRE of interest. For this, relevant ATAC-seq “peaks” were recovered using BEDOPS (3.1.1 v2.4.39; *--ec -element-of-1* parameters specified(58)). HOMER’s *findMotifsGenome*.*pl* program was then executed the recovered ATAC-seq “peaks”, using the same parameters and motifs threshold criteria as specified above.

### Clustering analyses

Clustering analyses on RNA-seq data were done using the *pheatmap* R package. TPM-normalized data was clustered according to z-scores across conditions for each differentially expressed gene. The optimal number of clusters (*k*) was determined using the *NbClust* package(59), which calculates multiple indices to determine *k*. For H3K27ac ChIP-seq data, the same process as above was done using z-scores from the *simpleNorm* datasets for each of the differentially enriched regions.

### Gene ontology analyses

Metascape with default parameters (https://metascape.org/) was used to perform gene ontology analyses on groups of genes of interest (RefSeq identifiers)(60). Terms associated with KEGG pathways were used unless otherwise specified(61).

### Data visualization

Plots were made in R using the *tidyverse, pheatmap, rcartocolor* and *ggpubr* packages. The UCSC genome browser was used to visualize ChIP-seq and ATAC-seq data(62).

### Immunolabelling

Mice were anesthetized with an intraperitoneal injection of a lethal dose of ketamine (91 mg/ml) and xylazine (9.1 mg/ml) and then rapidly perfused transcardially with 0.9% saline, followed by 4% paraformaldehyde (PFA) in 0.1 M borax buffer, pH 9.0 at 4 °C. After perfusion, brains were rapidly removed from the skulls, postfixed for 2 days, and then placed in a solution containing 20% sucrose diluted in 4% PFA/borax buffer (pH 9.0). The frozen brains were mounted on a microtome (Leica Biosystems), frozen with dry ice, and sliced into 30 μm coronal sections using sliding microtome. The slices were collected in a cold cryoprotective solution (0.05 M sodium phosphate buffer, pH 7.3, 30% ethylene glycol, 20% glycerol) and stored at −20°C.

Free-floating brain sections were washed and immunolabelled with rabbit anti-mouse ionized calcium binding adaptor 1 (Iba1) (1:1,000, Wako Chemicals, 011-27991) primary antibody overnight at 4 °C and revealed with a biotinylated goat anti-rabbit IgG secondary antibody (1:1,000, 2 hat RT, Vector Laboratories, VECTBA1000), the VECTASTAIN ABC Elite HRP kit (Vector Laboratories) and 0.125 mg/ml of DAB peroxidase substrate solution (Sigma-Aldrich, D5637).

### Microscopy analyses

Two microphotographs for each of the striatum (caudate putamen) and the primary somatosensory cortex were taken on adjacent brain slices, between Bregma 0.13 mm and 1.09 mm, and then imported to ImageJ(63) for quantification of Iba1^+^ cells. For this, a 0.73 mm x 0.73 mm box was first drawn over regions of interest and encompassed Iba1^+^ cells were then counted. Cell counts over the two slices for each region were then averaged. All analyses were performed blindly. Statistical significance was assessed using Welch’s T-Test, with alpha level set at 0.05.

## General

We thank Sophie Vachon for assistance with mouse colony management.

## Author contributions

AMX, FDG and DG conceived the study. AMX and FDG performed mice experiments, collected and analyzed data. S Belhocine isolated microglia. NB, S Bitarafan and AF collected data. SF performed experiments and analyzed flow cytometry data. SR provided intellectual input. DG generated sequencing libraries, analyzed data and wrote the manuscript with assistance from AMX and FDG. DG also supervised all aspects of the project.

## Supplementary data

**Figure S1.**
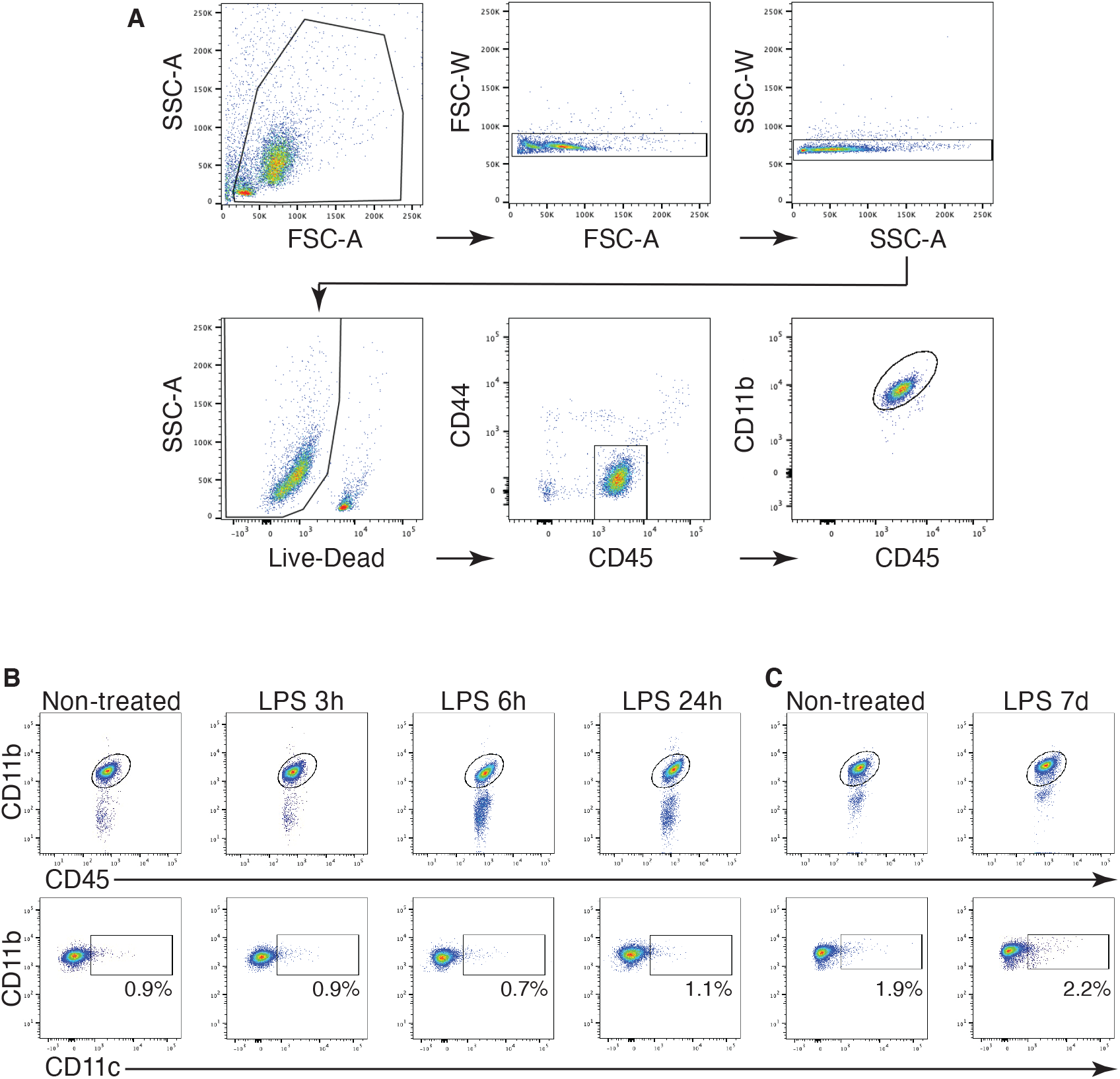
Gating strategy for isolation of microglia. **(A)** Microglia were defined as live-CD11b^+^CD45^+^CD44^Low^ cells. **(B, C)** Scatterplots of CD11b, CD45 and CD11c expression of microglia isolated from the brain of mice at different time points after LPS injection.

**Figure S2.**
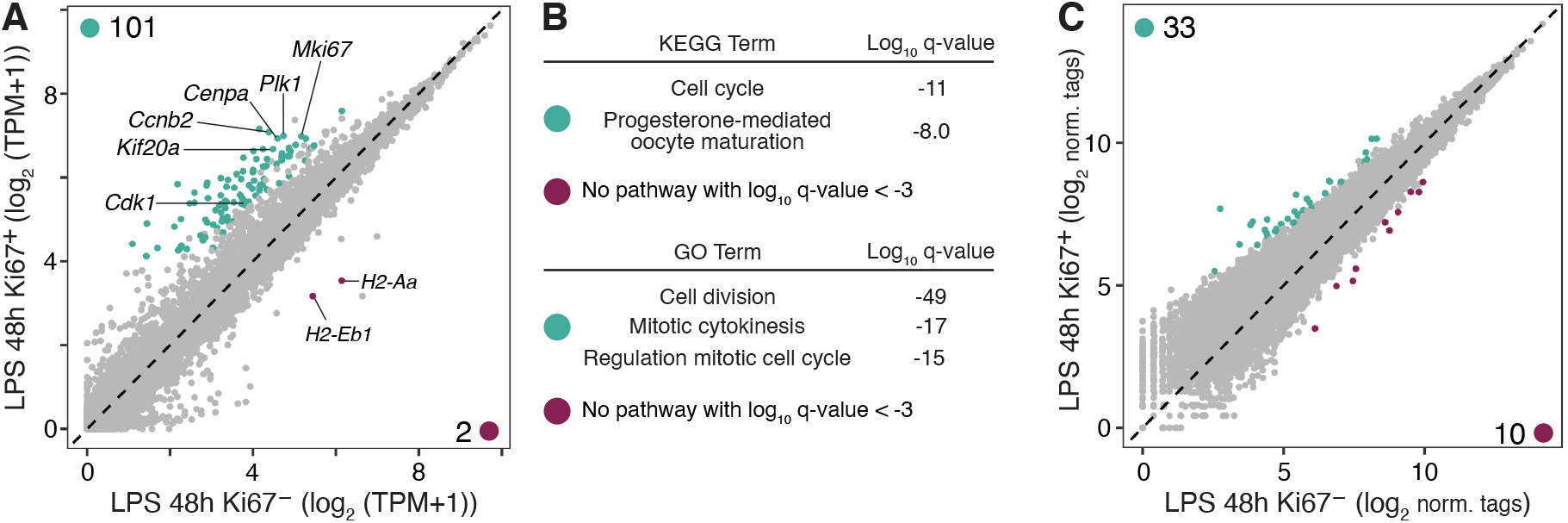
Gene and H3K27ac profiles of proliferative microglia isolated 48 h after LPS injection. **(A)** Scatterplot of RNA-seq data from RFP^High^/Ki67^**+**^ and RFP^Low^/Ki67^**−**^ microglia isolated from the brain of mice administered LPS 48 h earlier. Differentially expressed genes identified by DESeq2 (log2 FC ≥ 0.58, FDR-adjusted p-value ≤ 0.01) are colored coded. **(B)** KEGG pathways and Gene Ontology (GO) analyses of genes more highly expressed in LPS 48h Ki67^**+**^ vs LPS 48h Ki67^**−**^, and vice versa. **(C)** Scatterplot of H3K27ac abundance at distal CREs, comparing Ki67^**+**^ and Ki67^**−**^ from microglia isolated 48 h after LPS injection, as assessed by ChIP-seq. Differentially enriched distal CREs identified by DESeq2 (FDR-adjusted p-value ≤ 0.05, two-fold or greater tag difference) are colored coded.

**Figure S3.**
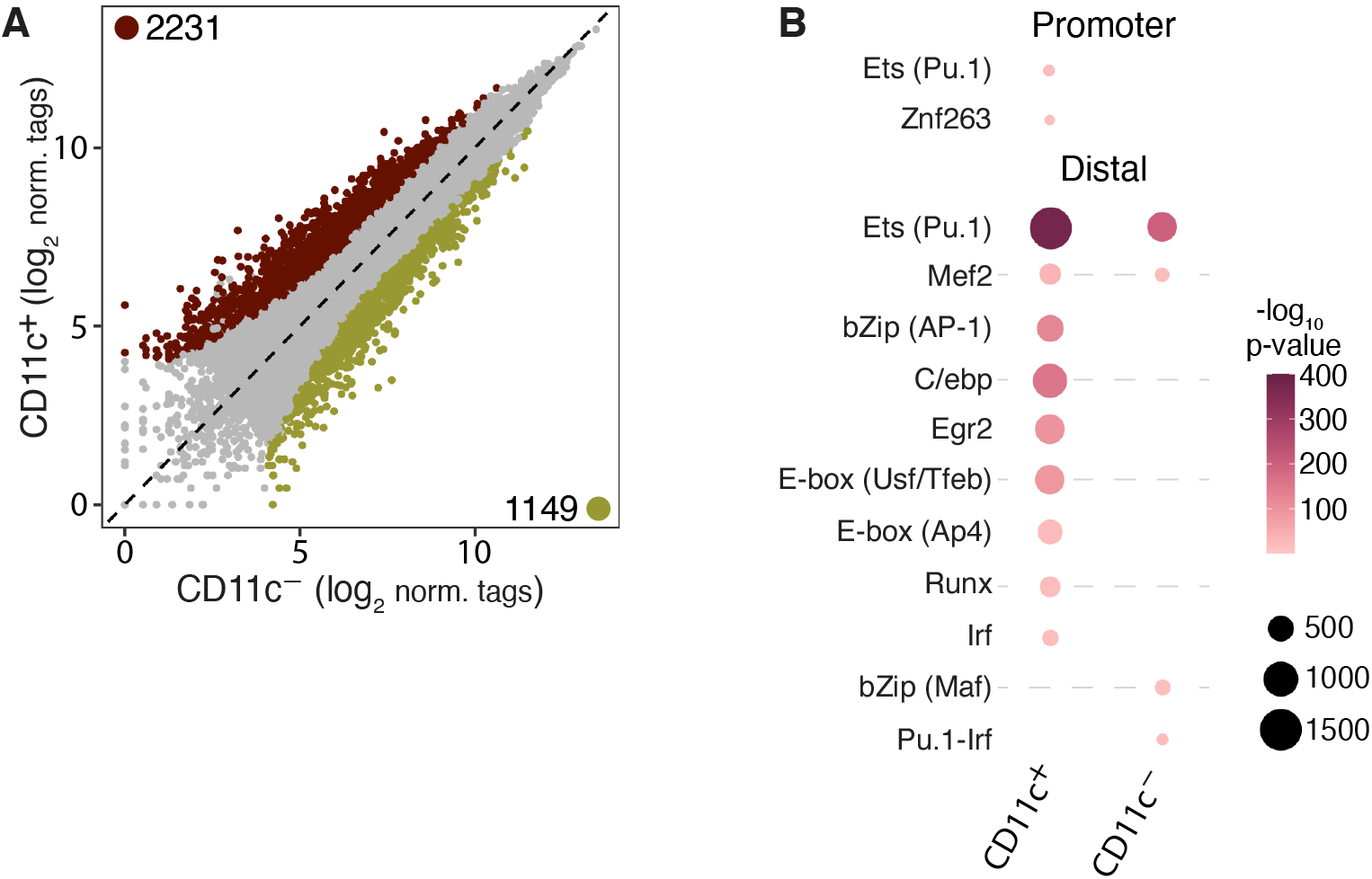
H3K27ac ChIP-seq data comparing CPZ CD11c^+^ and CPZ CD11c^−^. **(A)** Scatterplot of H3K27ac abundance at distal CREs. Differentially enriched CREs identified by DESeq2 (FDR-adjusted p-value ≤ 0.05, two-fold or greater tag difference) are colored coded. **(B)** Table summarizing results from de novo DNA motifs analyses performed on differentially active CREs identified in A.

### Tables

**Table S1. RNA-seq data associated with Figure 1**.

**Table S2. RNA-seq data associated with Figure 3**.

**Table S3. RNA-seq data associated with Figure 5**.

**Table S4. RNA-seq data associated with Figure 6**.

**Table S5. Global raw and TPM-normalized RNA-seq data used**.

